# Novel Methodology to Detect Physical Interaction of Inositol Phospholipid with Potassium Ion Channel Using Luminescence Resonance Energy Transfer

**DOI:** 10.1101/546283

**Authors:** Felix Chin, Ping-Chin Cheng

## Abstract

Membrane proteins like ion channels are located within a lipid environment. While the bulk lipids provide a supporting architecture for proteins, some lipids also play important signaling roles in modulating protein activity. This modulation can occur through direct interaction with proteins without involving second messenger pathways. Phosphoinositides are one such lipids, which have been implicated in interactions with a diverse range of proteins and such interactions mediate a variety of cellular functions. Towards understanding how these lipids play their roles in regulating protein functions It is necessary to detect interactions between proteins and lipids and to identify binding sites on proteins. However, detection of protein-lipid interactions has been difficult especially in their native environment because only a fraction of lipids are bound to proteins while the bulk is freely diffusing. The purpose of this study is to develop an experimental method capable of identifying protein-lipid interactions in membrane environments. The strategy involves luminescence energy transfer to locally illuminate lipids around proteins thereby suppressing the large background of light from unbound lipid molecules. The approach is applied to an inward rectifier potassium ion channel for its interactions with phosphoinositide lipids. Experiments show that it can successfully detect phosphoinositides bound to proteins, and when combined with mutagenic approaches, can further identify binding site. The approach thus provides a valuable methodology for probing protein-lipid interactions in a native-like lipid environment.

## Introduction

Membrane proteins play key roles in detecting and conveying signals across cell membranes. It is estimated that 20% of total human genes code for membrane proteins [1]. Ion channels are specialized integral membrane proteins that control cellular ionic homeostasis. By controlling membrane potentials of cells, they are involved in a diverse amount of physiological processes. For example, muscle cells depend on ion channels for their electrical excitability, nerve cells utilize channels to generate and propagate electrical pulses within neurons and across synapses, and somatosensory cells rely on ion channels to detect and transduce environmental cues including lights, sound and temperature. Dysfunctions of ion channels, due to inherited or acquired mutations, have been implicated to leading to a number of diseases. In the search for new drugs, ion channels are an important target.

Every ion channel responds to some specific activator(s). Aside from their activators, ion channels are also further regulated by cellular components. Such regulation can modify channel activity and dramatically alter the electrophysiological properties of both excitable and non-excitable cells. Many regulatory mechanisms exist, and the interaction with lipids represents one important type. Because ion channels physically resided in membranes, they are particularly prone to modulation by lipids. Many lipids are bioactive, especially those that are less abundant and their extent can be dynamically regulated. One classical example is phopshoinositides such as phosphatidylinositol 4,5-bisphosphate (PIP_2_). Signaling roles of the lipid as a precursor for the ubiquitous messenger molecules inositol triphosphate (IP_3_) and diacylglycerol (DAG) has long been appreciated. But as Hilgemann and his colleague discovered later [2], the lipid itself also directly interacts with ion channels and modulates their activity. Since then hundreds of publications have emerged showing that most ion channels are regulation targets of the lipid. In addition to PIP_2_, other lipids and their metabolic products such as cholesterol and sphingomyelin have also been implicated in regulation of ion channels.

Decades of study have now established lipid interactions as an important mechanism for modulation of ion channels. However, how the interactions occur in different ion channels and how they control channel activity have been less clear. In order to understand the problem, one critical step is to determine whether lipids directly bind to proteins and where they bind on the protein. Unfortunately, identifying protein-lipid interactions is notoriously difficult. The binding of lipids with proteins is energetically weak and the binding sites are often buried (in membranes). As a result, protein-lipid interactions cannot be analyzed by common protein-ligand binding assays. Instead, researchers have resorted to other means for studying such interactions [3]. One most popular approach is structural-functional studies where putative sites are mutated and the roles of the residues in binding are assessed based on whether the mutations affect lipid functions. The problem with this approach is that the gating of ion channels is a complex allosteric process. Residues may affect protein functions by altering gating properties rather than directly controlling binding. Indeed, work on inward rectifier potassium channels has resulted in numerous putative sites for phosphoinositide binding; however, most of these appear to affect gating only. In other approaches, isolated protein fragments containing putative binding sites or more completely, the entire protein, is analyzed for lipid interactions by immunochemistry methods in a way similar to western blot analysis. Thus these approaches have similar limitations as western blot analysis such as dependence on antibody specificity. More importantly the isolation or solubilization process may disrupt native binding sites. In rare cases, X-ray diffraction has been used to solve atomic structures of proteins bound with lipids; however, crystallization of membrane proteins is a daunting task, when not even mentioning the protein-lipid complex. So far, only structures of Kir2.2 and Kir3.2 in complex with PIP2 have been resolved at the atomic level [4, 5].

The lack of a general binding method capable of detecting protein-lipid interactions in their native environment has been one major factor impeding the study of lipid regulatory mechanisms of ion channels. This project was undertaken to address the problem. The strategy involves using fluorescence energy transfer (FRET) to directly detect protein-lipid interactions. To limit background fluorescence from non-bound lipids, a modified FRET, luminescence energy transfer (LRET) was used. Application of the approach to a prototype lipid-interacting ion channel led to successful detection of lipid interactions and identification of binding sites, thus demonstrating experimental feasibility of the approach.

## Materials and Methods

### Protein Expression and Purification

The full-length cDNA encoding human Kir2.1 or its mutants was subcloned into a FastBac1 expression vector. A cassette sequence containing a maltose binding protein (MBP) followed by a ~20 amino acid linker containing a consensus sequence for thrombin cleavage site was introduced at the N terminus of the protein. A lanthanide binding tag (LBT) was introduced into the channel at the turret position by replacing residues SKEGK with the LBT sequence: YIDTNNDGWYEGDELLA. The construct was transformed into E.coli DH10Bac strain, and the resultant bacmid was isolated and transfected into SF9 cells to make recombinant baculovirus. Sf9 cells in suspension culture were infected with recombinant virus and harvested 72 hours post-infection. Cells were lysed by nitrogen cavitation (Parr bomb). The lysates were collected and centrifuged at 800g for 20 min to remove nuclei and intact cells. The membrane fraction was collected by ultracentrifugation at 100,000 g for 1 hour and re-suspended in Buffer A (200 mM NaCl, 2 mM TCEP, 10% glycerol, 50 mM HEPES, pH 8.0). Membranes were solubilized by addition of 1% DDM in the same buffer. After incubation for 20-30 min at 4°C, soluble and insoluble fractions were separated by centrifugation at 18000 g for 15 min. The soluble fraction was loaded onto a column packed with 1.5-2 ml amylose resins at a flow rate of 0.2 ml/min. The resin was washed with 20 column volumes of wash buffer (Buffer A) containing 0.5 mM DDM and 10 μg/ml lipid mixture. Elution was done in the same buffer in the presence of 20 mM maltose. All steps were carried out at 4°C or on ice. Size-exclusion chromatography (SEC) was performed on a superpose 6 column (10mm/300mm), using Buffer A as the running buffer. Samples from the affinity chromatography were used either directly or submitted to an additional SEC step to collect tetramer peak fractions. Proteins were concentrated in an Amicon Ultra concentrator tube (100 kDa molecular weight cutoff, Millipore). Final protein concentration was quantified based on tryptophan fluorescence measured at 340 nm on a fluorometer (Hitachi).

### Fluorescence Spectroscopy

Time resolved luminescence was measured on a cuvette-based fluorescence spectrometer. Short excitation light pulses at 285 nm were formed by passing light from a Xenon lamp through a mechanical chopper operating at 20 kHz. Emission lights were detected after excitation light pulses to avoid prompt fluorescence. Donor emission was collected at 490 nm and acceptor sensitized emission at 510 nm. The background signal was monitored before every experiment to ensure it was negligible. The LRET lifetimes were fit by either single or two exponentials with Origin software (OriginLab Corp).

### Liposome Reconstitution

Proteoliposomes were prepared using the detergent-mediated reconstitution method according to [6]. Different lipids (POPC/POPE/POPG), stored in chloroform, were mixed at a ratio of 2:1:1, which emulates the composition of soybean lipids. The mixture was dried under nitrogen stream, and residual chloroform was further evaporated under vacuum for ~2 hours. A total of 250 µg lipid mixture was used for each reconstitution. The dried lipidic film was rehydrated in 50 µl reconstitution buffer (5mM MOPS, 200mM KCl, pH 7) containing 2 mM TCEP. Liposomes were formed and homogenized by alternately freezing and thawing the hydrated lipids in liquid nitrogen and warm water for 6-10 cycles, followed by bath sonication for ~2 minutes. Fluorescent lipids (Bodipy-FL-PIP2 from Echelon and Bodipy-FL PC from Life Technologies), when needed, were added to the basic lipid mixture at appropriate ratios during liposome preparation.

For reconstitution, 50-100 µl of preformed liposomes (5mg/ml) were equilibrated with DDM (final concentration 4.8 mM) at room temperature for 1 hour. Purified proteins were then added to the lipid/detergent mixture (1:10-20 protein: lipid ratio), and the ternary protein/lipid/detergent mixture was incubated at room temperature for 1 hour. Polystyrene beads (Bio-Beads SM2, Bio-Rad) were then added at a wet weight of 3-5 mg to remove detergents and the mixture was incubated at room temperature for 1 hour. This Bio-Bead treatment was repeated for three more times, and for the last one, the mixture was incubated overnight at 4 ^°^C on a shaker. Proteoliposomes were collected by ultracentrifugation of the sample for 1 hour at 4 ^°^C. The pellet was re-suspended in reconstitution buffer and flash-frozen in aliquots for storage at −80 ^°^C.

### Proteoliposome Electrophysiology

Multilamellar proteoliposomes suitable for patch-clamp recording were formed following protocols in [7]. Frozen proteoliposomes (4–8 μl) were thawed and supplemented with sucrose to a final concentration of 15-20 mM. The mixture was transferred to a glass coverslip for dehydration in a vacuum desicator for ~2 hours at room temperature. The dried lipidic film was then rehydrated by addition of 10 μl reconstitution buffer followed by incubation overnight at 4°C in a humidified chamber. A small amount of rehydrated proteoliposomes was used for each patch clamp experiment. Currents were recorded with an Axopatch 200B amplifier (Molecular Devices).

## Results and Discussions

### The Theory

#### 1. FRET versus LRET

Fluorescence resonance energy transfer (FRET) has been widely used for detecting molecular interactions. In this approach, two molecules are conjugated respectively with two different fluorophores, one as the donor and the other as the acceptor. Upon excitation the donor transfers its energy to a neighboring acceptor by non-radiative dipole-dipole interaction. The efficiency of this energy transfer is inversely proportional to the sixth power of the distance between the donor and the acceptor. Due to this sharp distance dependence, FRET occurs efficiently within a proximity of 50-100A, which is the range of inter-and intra-molecular distances and thus makes the approach particularly suitable for probing molecular interactions.

However, one important limitation for application of FRET in practice is unknown or variable donor-to-acceptor stoichiometry. Unpaired donors and/or acceptors cause background fluorescence due to direct excitation. In the extreme case where one partner is in excess concentration, direct fluorophore emission outweighs FRET signal, so that detection of a small level of FRET against a large background of fluorescent labels that are not undergoing FRET becomes technically prohibitive. This is exactly a problem occurring with protein-lipid interactions. The lipids, even the least abundant ones in the membrane, far exceed proteins in amount. Because of this problem the standard FRET is not readily useful for detection of protein-lipid interactions.

One potential solution to the problem is luminescence energy transfer (LRET), a modified FRET [8]. LRET is based on the same principles as FRET, but instead of using two organic fluorophores, uses a lanthanide donor and an organic acceptor. Lanthanides differ from organic fluorophores in terms of lifetimes in the excited state. While the excited state of organic fluorophores lasts only nanoseconds, the excited state of lanthanides have millisecond lifetimes. This orders-of-magnitude lifetime difference makes luminescence and fluorescence temporarily separable. Thus in the LRET approach a pulse of light is employed to excite the donor-acceptor sample and detection of emission starts after a few tens of microsecond delay. All prompt fluorescence including auto-fluorescence and direct acceptor emission are quenched before detection starts so they will not contribute to measurement. Furthermore, lanthanides have sharply spiked emission spectra. Acceptor fluorophores can be properly chosen so that only acceptor emission, not donor emission, is detected. For example, terbium is dark at wavelengths where fluorescein and tetramethylrhodamin emit. Owing to this spectra distinction, sensitized acceptor emission from only donor-acceptor pairs becomes possible to measure. Its lifetime follows the lifetime of the donor, but without contribution from donor-only species. Therefore by both temporal and spectra discriminations LRET is capable of eliminating contaminating background due to either direct acceptor fluorescence or donor emission. These advantages make LRET potentially useful for detecting energy transfer in complex labeling mixtures such as proteins and lipids in membranes.

#### 2. Feasibility of LRET for detection of protein-lipid interactions

An additional issue that may limit the applicability of LRET for detection of protein-lipid interactions is nonspecific energy transfer. The lipids in the membrane may be so crowded that non-bound lipids are also in proximity of proteins within the distance of energy transfer. In this case, the energy transfer will consist of two pools, one from the donor to specifically bound lipids and the other from the donor to freely diffusible ones. Conceivably, at very high lipid concentrations, the nonspecific transfer may overwhelm the specific transfer, limiting detectability of interacting lipids. The feasibility of LRET for detecting protein-lipid interactions will thus depend on lipid concentrations.

Does LRET have an adequate resolution at a *working* lipid concentration? Fortunately, signaling lipids tend to have low abundance in membranes. PIP_2_, for example, account for ~1% of total membrane lipids. Among them a significant portion is further sequestered leaving free ones at even lower levels. At this concentration (1%), there is one PIP_2_ molecule every 100 lipid molecules. Each lipid molecule has a surface area around 70Å^2^. Thus one PIP_2_ molecule is present over an area 7000 Å^2^. Assume that an ion channel has a radius of 45 Å [9]. The nearest PIP_2_ from the protein will occur at a distance *d* = (7000/π +45^2^)^1/2^ = 65 Å. Since energy transfer decreases sharply with distance, it can be assumed that energy transfer to non-interacting lipids is dominated by the first shell of non-interacting lipids. The rate of this energy transfer follows *k*_*nt*_ = τ_D_^-1^(*R*_0_/*R*)^6^ where τ_D_ is the time constant of the donor lifetime (in the absence of acceptor), *R*_0_ is the characteristic distance at which half of the energy is transferred, and *R* is the donor-acceptor distance. With typical values *R*_0_=44 Å (fluorescein) and τ_D_ ~2 ms, the rate of energy transfer non-interacting lipids *k*_*nt*_ is estimated at 47 s^-1^. On the other hand, lipids bound to proteins have a shorter distance between donor and acceptor. According to crystal structures of Kir channels, the PIP2 molecule is located about 44 Å from the turret (where the donor is located as described below). At this distance the rate of energy transfer to bound lipids *k*_*st*_ is ~500 s^-1^. The energy transfer to bound lipids has an efficiency >10 times to non-bound ones. The actual efficiency will be even higher if long acyl chain PIP_2_ is used. LRET is therefore dominated by interacting lipids.

In general, the distance between the protein and the first-shell non-interacting lipids can be derived as *d* = (*d*_*0*_^2^+70/π*C*)^1^/^2^ where *d*_0_ is the radius of protein and *C* is lipid concentration in percentage. This equation suggests that the distance is related to the square root of the concentration and thus has a relatively slow dependence on concentration changes. It is noted that applicability of LRET does not require the bound lipid to proteins to dominate energy transfer. LRET measures the donor lifetime, which is given by τ = (*k*_nt_+*k*_*st*_+*k*_D_)^-1^ where *k*_*D*_=1/τ_*D*_ is the rate of decay in the absence of acceptor, and *k*_nt_ and *k*_*st*_ are as described above. As long as *k*_*st*_ is not negligible as compared to *k*_*nt*_, its contribution to the lifetime of the donor should remain detectable. Thus the theory predicts that LRET is applicable to detection of protein-lipid interactions at *working* lipid concentrations.

### Experimental Results

To validate its experimental applicability I have applied the LRET approach to probe interactions of ion channels with lipids. In particular, I have focused on an inward rectifier potassium channel (Kir2.1, Fig.1A), a prototype lipid-interacting ion channel that was among the first found to be directly regulated by phosphoinositides such as PIP_2_. This channel was chosen because there is a wealth of information on the channel, which can be explored for crosscheck of findings. However, despite extensive research, there are still issues and controversies on interactions of the channel with phosphoinositides, such as how many binding sites exist and where they are located on protein. Application of the LRET approach will provide independent measurements to address these issues.

**FIGURE 1.**
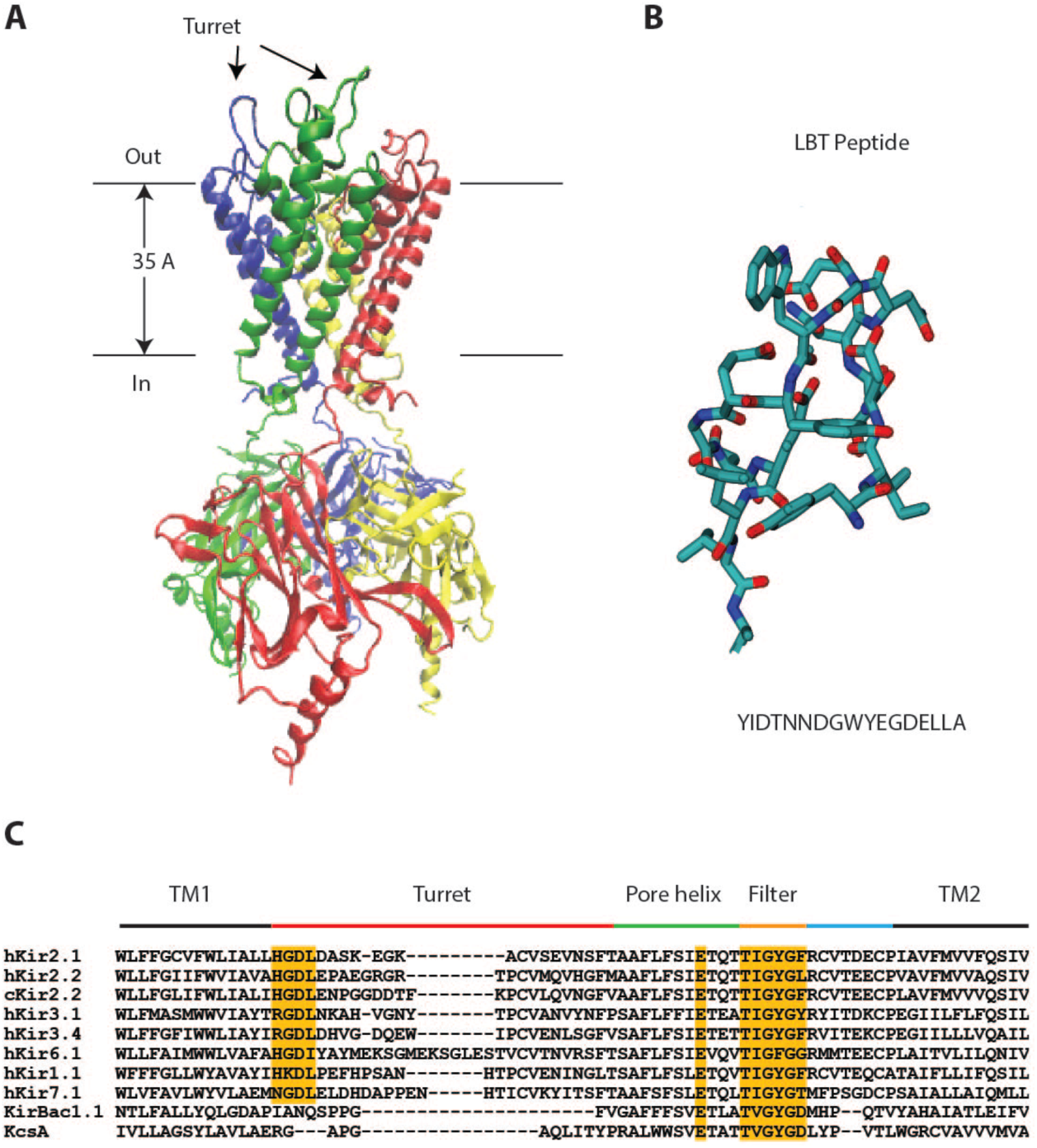
Incorporation of lanthanide binding tag (LBT). (A) Atomic structure of human Kir2.1. The channel is a homotetramer with each subunit containing two transmembrane segments (TM1 and TM2) and a pore loop between them. (B) Structure of a genetic LBT. The amino acid sequence of the tag is shown below the structure. (C) Sequence alignment of Kir channels in the external pore region, including human Kir2.1 (GI: 8132301), human Kir2.2 (GI: 23110982), chicken Kir2.2 (GI: 118097849), human Kir1.1 (GI: 1352479), human Kir3.1 (GI: 1352482), human Kir3.4 (GI: 1352484), human Kir6.1 (GI: 2493600), human Kir7.1 (GI: 3150184), KirBac1.1 (GI: 33357898), and KcsA (GI: 39654804). The alignment shows that the turret loop is least conserved. Both its amino acid composition and size vary considerably among different members, suggesting that the structure of the region is likely flexible. The LBT sequence was inserted in human Kir 2.1 in place of the SKEGK sequence before the gap.

#### 1. Incorporation of Lanthanide Binding Tag (LBT)

A major challenge in application of the LRET approach is introduction of a lanthanide binding tag into subject protein. Because lanthanides have a low extinction coefficient, these chelating molecules or motifs also serve as sensitizer antennae to excite lanthanides. Two types of binding tags are available, either an organic chelate molecule or a genetic binding motif. The organic chelate is incorporated into the protein by labeling cysteine residues using thio-reactive reagents. The labeling has disadvantages such as incomplete or nonspecific labeling. More importantly it requires removal of endogenous cysteine residues in proteins, which may not be feasible for large proteins such as ion channels containing a large number of cysteine residues. For these considerations I have chosen to use a genetic binding tag introduced by Imperiali and collaborators [10]. This tag consists of 17 amino acids (YIDTNNDGWYEGDELLA) arranged in a structure similar to EF hand Ca^2+^-binding motif but has a high affinity and specificity for terbium ions (Tb^3+^) (Fig.1B). The two termini of the motif are also closely spaced, so it can be incorporated into proteins with minimal distortion on protein structures. The tryptophan (Trp) and tyrosine (Tyr) residues within the tag are close to the bound lanthanide and serve as sensitizers to transfer energy to the bound lanthanide when illuminated by ultraviolet (UV) light.

Ideally, the LBT would be best placed in protein at a position closest to the lipid binding site. That way the energy transfer from the LBT donor to the bound lipid acceptor is maximized. On the other hand, the tag needs to be outside (on extracellular domains) in order to be able to titrate Tb^3+^. For Kir channels which have only two transmembrane segments per subunit (Fig.1A), this means that the LBT has to be introduced at the extracellular pore loop region. This region consists of a turret domain followed by a pore helix, a selectivity filter and then a linker domain to the second transmembrane helix. Sequence alignment of Kir homologs indicate that the turret region is least conserved and has a variable length (Fig.1C). The divergence of sequence in this region suggests that it is not functionally essential, making it a good candidate place for insertion of LBT.

By mutagenesis an oligonucleotide sequence encoding the LBT was designed and inserted into the open reading frame of the Kir2.1 channel. The LBT was introduced in the turret loop in place of the SKEGK fragment at the gap (Fig.1C). The resulting chimeric Kir2.1-LBT channel was expressed in mammalian cell lines (HEK293) for examination of functions. Large currents comparable to wild type channels were recorded from these channels on the cell surface (Fig.2A). The chimeric channel was also well expressed in SF9 cells for production of recombinant proteins. Importantly, they remained stable to survive purification, and the purified proteins could be functionally reconstituted into artificial liposomes made of synthetic lipids containing PIP_2_ (Fig.2B). Thus the insertion of the LBT did not distort channel functionality. Since Kir channels require PIP_2_ for function, the responsiveness of the LBT chimera also implies that the lipid binding site remained intact.

**FIGURE 2.**
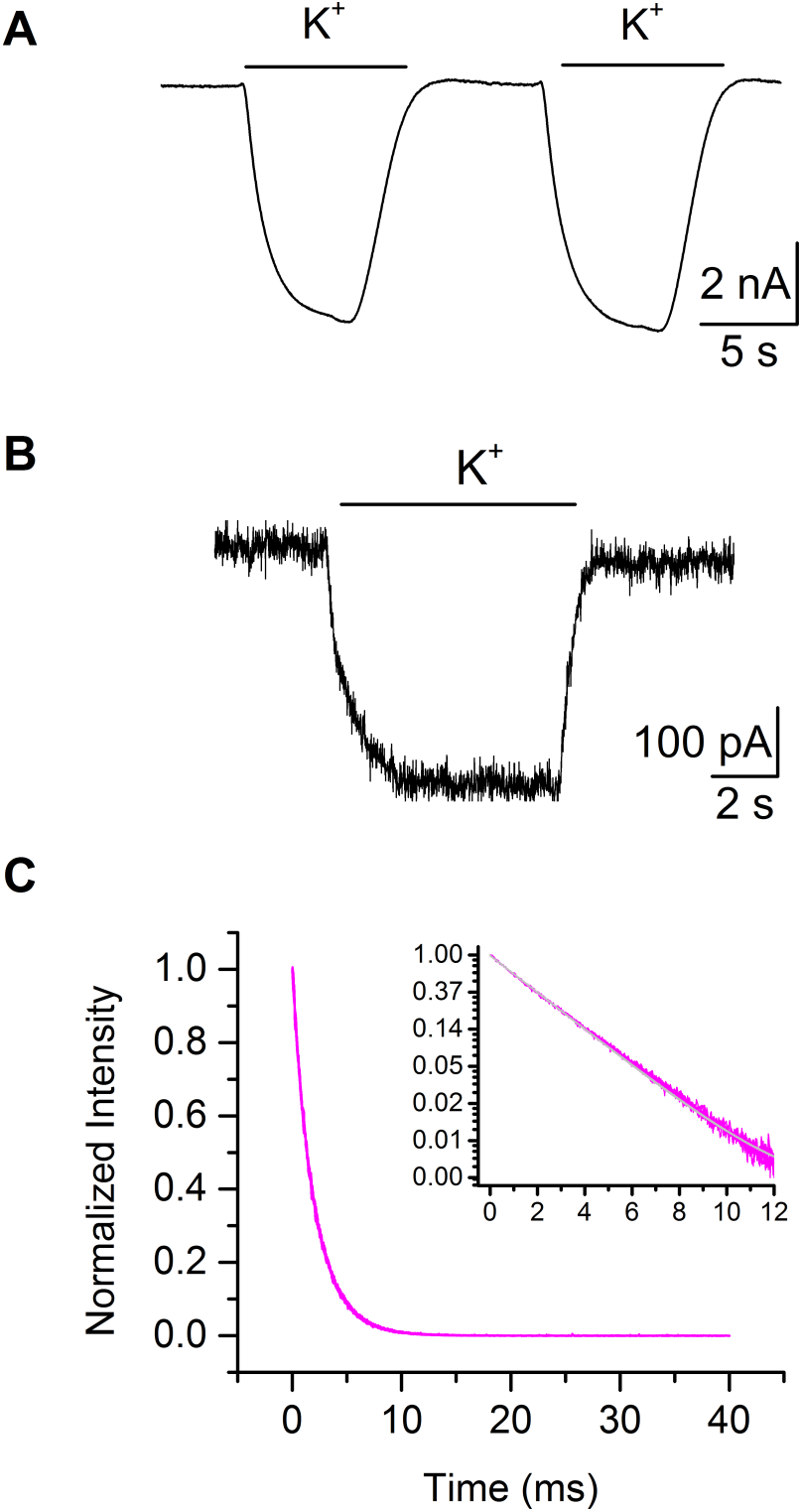
Phenotypes of the LBT-containing chimeric channel. (A) Whole-cell currents of the chimeric channel expressed in SF9 cells. The current was evoked by perfusion with a saline solution containing 150 mM potassium ions. (B) Macroscopic current from reconstituted chimeric channels in liposomes. Holding potential was −60 mV at both conditions. Control cells or liposomes without channels showed no detectible activity (data not shown). (C) Tb^3+^ luminescence measurement from reconstituted proteoliposomes containing the chimeric channel. The inset shows the decay in the logarithm scale. The time course of the decay was fit by a single exponential with time constant 2.01±0.01 ms (n=4). For B and C, the artificial lipids (POPE/POPC/POPG) were supplemented by 0.1% nonfluorescent PIP_2_.

The inserted LBT was also kept intact and functional. As a test, the donor-only emission was measured in the absence of acceptor molecules. Fig.2C illustrates time-resolved Tb^3+^ luminescence measured at 490 nm following a brief excitation at 285 nm. The emission is Tb^3+^-specific; priori to addition of Tb^3+^ ions, no emission was detectable. The emission also decayed in an exponential manner characteristic to free LBT in solution [10]. The decay has a monoexponential time course with a time constant of 2.01 ± 0.01 ms (n=4), which is slightly shorter than but comparable to that of free LBT in solution. Terbium ions at a concentration of a few micromolars appear adequate to emit a strong luminescence signal. Since ion channels are tetramers, a higher concentration in the range of 10-20 micromolars was used instead in order to fully saturate Tb^3+^ binding on multiple channel subunits.

Of note, the structure of the pore domain of Kir channels is representative to a large number of ion channels. These include the six-transmembrane segment family of voltage-gated ion channels and the superfamily of transient receptor potential (TRP) channels. Thus, insertion of LBT into the turret region provides a general strategy to incorporate LBT into other pore loop-containing channels, thus making it possible to apply LRET to investigate lipid regulations of those channels as well.

#### 2. Interaction with Phosphoinositides

For detection of interaction with phosphoinositide lipids, purified Kir2.1-LBT channels were reconstituted into chemically defined liposomes composing of three basic lipids POPC/POPE/POPG mixed at a 2:1:1 ratio and supplemented with a fluorescent PIP_2_ at a functionally required level (0.1%). Patch-clamp recording experiments confirmed functionality of reconstituted channels in this membrane environment. The fluorescent PIP_2_ was conjugated with Bodipy-FL fluorophore on one of its acyl chains. This fluorophore had an absorption peak around 490 nm so it could be paired to the Tb^3+^ donor for efficient energy transfer. Its emission was around 520 nm where Tb^3+^ did not luminesce, so that sensitized acceptor emission could be measured from only donor-acceptor pairs.

Fig.3A illustrates the time-resolved measurement of sensitized acceptor emission. The decay followed a bi-exponential time course with a main component at time constant of 0.82±0.03 ms (n=4), which was identified to arise from LBT-bound Tb^3+^ luminescence. The other component had a minor contribution (~20% in amplitude), with time constant τ=1.53 ± 0.15 ms (n=4), which may account for contributions from contaminating sources such as non-bound PIP_2_, instrument noise and nonspecifically bound Tb^3+^ by proteins and lipids.

**FIGURE 3.**
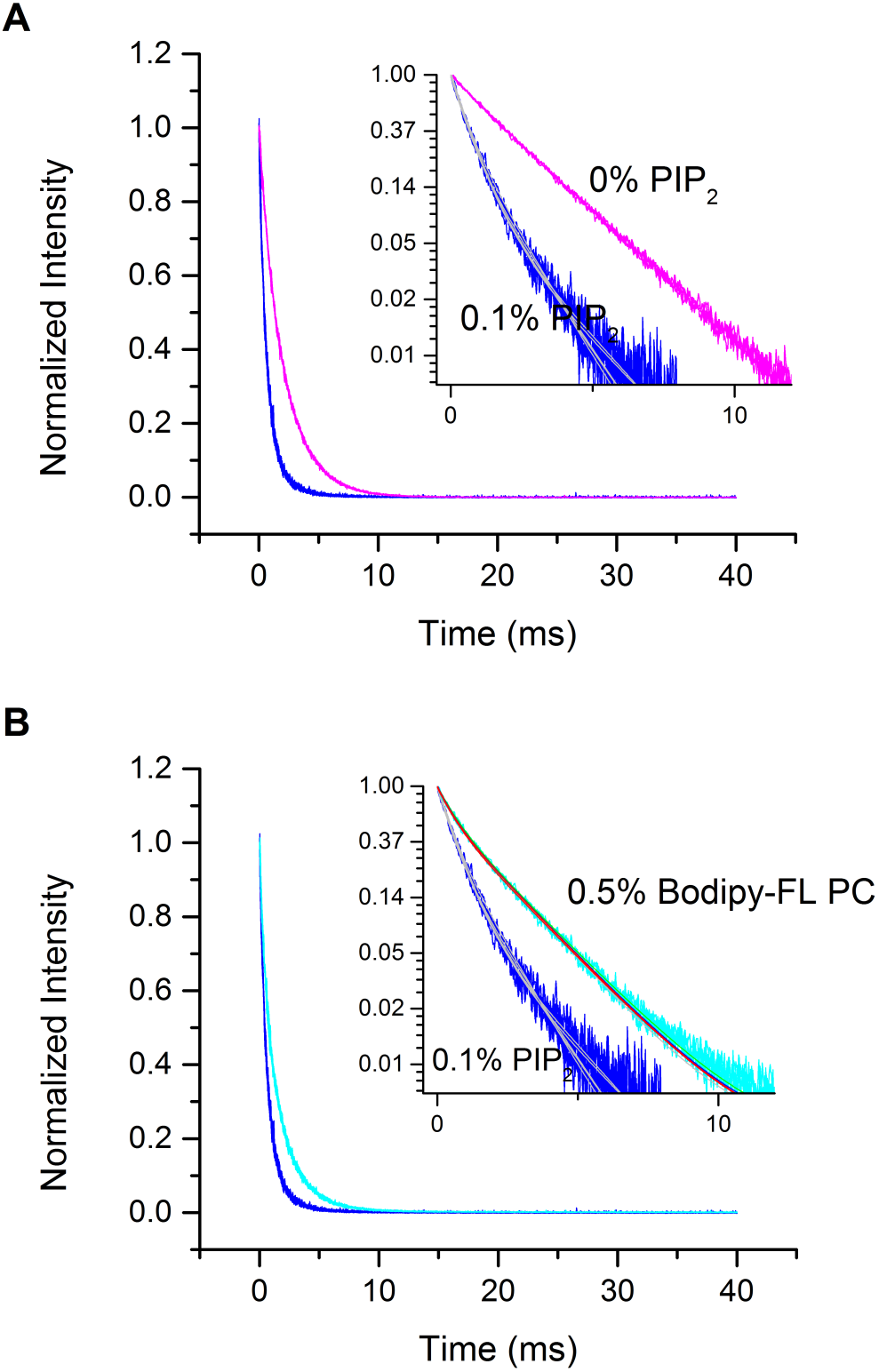
Detection of phosphoinositide binding. (A) Tb^3+^ luminescence decay in the presence of 0.1% Bodipy-FL PIP2. Also shown is the control time course in the absence of fluorescent PIP_2_. The inset shows the decay in the logarithmic scale. When fluorescent PIP_2_ was added, Tb^3+^ luminescence decay became considerably faster. The time course was fit by two exponentials with the major component having a short time constant 0.82±0.03 ms (n=4). This lifetime corresponds to a donor-acceptor distance ~37 Å. (B) Contribution of non-bound lipids. Bodipy-FL phosphatidylcholine (PC), which presumably does not bind to the channel, was used to mimic non-bound acceptors. Even at a concentration five time that of PIP_2_, Bodipy-FL PC resulted in only moderate acceleration in Tb^3+^ luminescence decay. Exponential fitting of the decay gave a major component with time constant 1.89±0.02 ms (n=5). Energy transfer in the presence of PIP_2_ was therefore dominated by bound PIP_2_ molecules.

Compared to the donor-only lifetime measurement, the presence of PIP_2_ accelerated Tb^3+^ luminescence decay by nearly three folds, suggesting an efficient energy transfer from the donor to the acceptor. The efficiency of the energy transfer is given by *E* = 1 - τ/τ_D_, which is estimated about 60%. With a characteristic distance *R*_0_ = 39 Å for the Tb^3+^ - Bodipy-FL donor-acceptor pair [11], this energy transfer efficiency corresponds to a donor-acceptor distance about 37 Å. X-ray crystal structures of Kir channels placed the distance between the turret and the PIP_2_ binding site at about 44 Å. The difference in the two distances probably had occurred for two reasons. One is that the fluorophore was conjugated at the end of acyl chains. The other is that the Bodipy-FL PIP_2_ used for LRET measurement was also long acyl chain PIP_2_, which brought the fluorophore further closer to the turret. Despite these considerations, the LRET measurement corresponds well to the distance obtained from crystal structures of these channels. This confirms the reliability of LRET measurement.

To further corroborate that the detected energy transfer was contributed by PIP_2_ molecules bound to channel proteins, the same experiment was repeated using a fluorescent phosphatidylcholine (PC) lipid instead of PIP_2_. The PC lipid, which has a neutral head group, presumably does not bind to the site where PIP_2_ binds. Thus the experiment provides a control reference for energy transfer to non-bound lipids. Fig.3B shows the sensitized acceptor emission in the presence of 0.5% Bodipy-FL PC. Notice that this concentration was five times that of Bodipy-FL PIP_2_ used in the previous experiment. The Tb^3+^ luminescence decay involved a major exponential component with time constant 1.89±0.02 ms (n=5). This time course was slightly faster than with donor-only, indicating that energy transfer did occur to non-bound lipids. However, the extent of the acceleration caused by Bodipy-FL PC, albeit at a higher concentration, is secondary compared to the acceleration caused by Bodipy-FL PIP_2_. Thus the strong LRET observed in the presence of PIP_2_ cannot be accounted for by energy transfer to neighboring diffusible PIP_2_ molecules around channel proteins; instead it must result from PIP_2_ molecules specifically bound to channel proteins.

#### 3. Interaction Site for PIP_2_

While PIP_2_ is known to interact with Kir channels, the location and structure of the binding site has been less certain. Electrophysiological or biochemical assays have implicated many residues for PIP_2_ binding, such as H53, R67, R82, K182, K185, K187, K188, R189, R218, K219, K228 and R312 [12-15], which are located throughout the N-and C-termini. Mutations of these residues are able to abolish PIP regulation and activation of the channels. On the other hand, crystal structures of Kir2.2 and Kir3.2 channels bound to PIP_2_ show only one specific site per subunit [4, 16]. This result was further elucidated with a PIP strip assay [17], which suggests R218 as one most essential residue for PIP_2_ binding. However, these later approaches including X-ray crystallography used proteins solubilized by detergents. Thus questions remain whether the discrepancy on PIP_2_ binding site is due to use of different experimental assays. Conceivably, solubilization of the channel by detergents may disrupt native binding sites that were identified by functional assays.

The LRET approach, which is capable of detecting direct binding in native-like membranes, provides an opportunity to potentially resolve the problem. If a residue is important for lipid binding, mutation of the residue will disrupt the binding site. The loss of specifically bound lipid will then result in reduction in luminescence energy transfer, and this change should be experimentally detectable. For illustration, the two critical residues R218/R219 as evident in crystal structures and reinforced by [17] were neutralized by substitution with glutamine. Functional studies by patch clamp recording showed that these mutations almost completely abolished channel activation as if PIP_2_ was no longer able to bind to the channel (data not shown). To “see” if PIP_2_ was indeed lost from the channel, the mutant channel (Kir2.1-LBT-R218/R219) was purified and reconstituted into liposomes containing 0.5% Bodipy-FL PIP_2_. This PIP_2_ concentration was five times that used for the wild-type-like Kir2.1-LBT channel. However, despite the increase in the PIP_2_ concentration, the decay of Tb^3+^ luminescence became much slower than previously observed with the wild-type-like Kir2.1-LBT channel in the presence of 0.1% PIP2 (Fig.4A). The decay had a time constant τ ~ 1.86±0.03 ms (n=5). A slower decay indicates less energy transfer, consistent with the notion that the mutations of R218Q/R219Q disrupted PIP2 binding.

**FIGURE 4.**
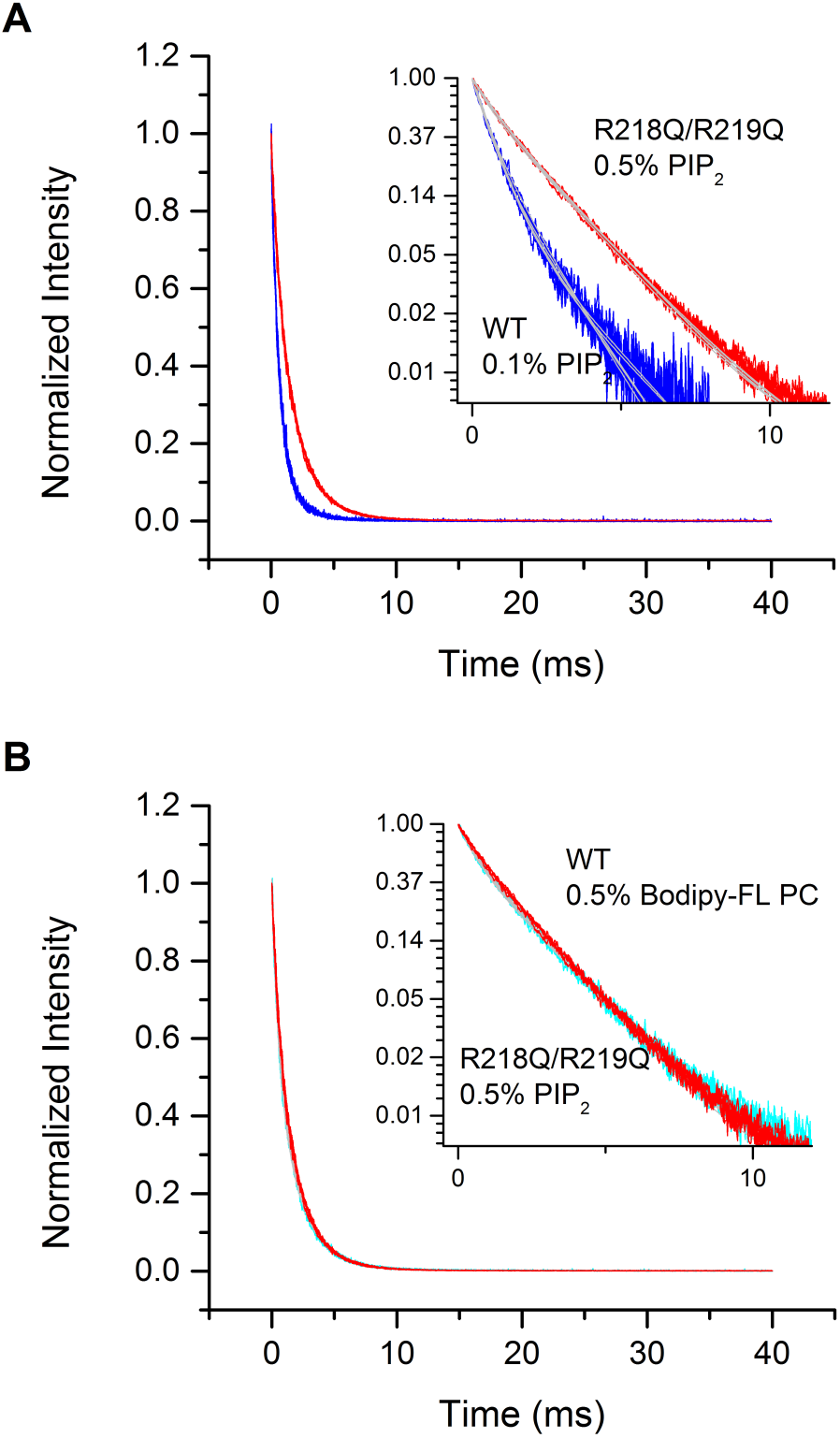
Identification of PIP_2_ binding site. (A) R218/R219 mediate PIP_2_ binding. Neutralization of the two residues considerably slowed down Tb^3+^ luminescence decay, as compared to the “wild-type” Kir2.1-LBT (in the presence of 0.1% Bodipy-FL PIP_2_). Since the PIP_2_ concentration was actually higher (0.5%) for the mutant (which would result in more nonspecific energy transfer), this reduction in the energy transfer is indicative of loss of bound lipids. (B) R218/R219 is the sole PIP_2_ binding site. The time course of Tb^3+^ luminescence decay for the R218Q/R219Q mutant (in the presence of Bodipy-FL PIP_2_) was superimposable with that of Kir2.1-LBT (in the presence of Bodipy-FL PC) when the two fluorescent lipid concentrations were kept the same. This implies a complete loss of PIP_2_ binding to the R218Q/R219Q mutant channel.

Are R218/R219 the sole binding site for PIP_2_ in Kir channels? This question was further assessed by comparing the energy transfer in the presence of PIP_2_ with the energy transfer in the presence of a control lipid that does not bind to the channel. For the latter case, energy transfer to 0.5% Bodipy-FL PC has an efficiency E ~ 6%. On the other hand, for a time constant 1.86 ms, the energy transfer for the mutant channel (Kir2.1-LBT-R218Q/R219Q) in the presence of 0.5% PIP_2_ was about 7%. The energy transfer efficiency in the two cases is nearly identical. Indeed, the two time courses of Tb^3+^ luminescence decay were superimposable. Since the energy transfer with Bodipy-FL PC arises from diffusible acceptors, the energy transfer for the mutant channel (Kir2.1-LBT-R218Q/R219Q) likely does not involve specifically bound PIP_2_ either. The LRET measurement thus supports that mutations of R218Q/R219Q are adequate to abolish PIP_2_ binding entirely and that Kir channels contain only a single PIP_2_ binding site at each subunit. Other residues that have been implicated by functional studies likely affect phosphoinositide sensitivity by mediating the gating of the channel rather than controlling PIP_2_ binding.

## Summary

Towards understanding how proteins are regulated by lipids it is necessary to identify the location and structure of lipid interaction sites. Methods for detection of molecular interactions are many, but few are applicable to protein-lipid interactions. The problem occurs because unbound lipids are in excess concentration creating a large background. The LRET approach is capable of detecting donor-acceptor interactions on large background fluorescence. Both theoretical and experimental results are presented to demonstrate feasibility of the approach for detection of protein-lipid interactions. When combined with mutagenic experiments it also allows for determination of the molecular basis of the interaction. For the first time, the approach was able to directly detect phosphoinositide interactions with ion channels in a native-like membrane environment. The results support that Kir2.1 channels contain a single binding site for PIP_2_ located at R218/R219, which agrees well with crystal structures of these channels and thus confirms the reliability of the approach. Although the present study has focused on the Kir2.1 channel, the same strategy was applicable to other membrane proteins especially ion channels containing similar pore domains, for which lanthanide binding tag can be similarly incorporated.

